# The *Staphylococcus aureus* regulatory program in a human skin-like environment

**DOI:** 10.1101/2023.10.24.563767

**Authors:** Flavia G. Costa, Krista B. Mills, Heidi A. Crosby, Alexander R. Horswill

## Abstract

*Staphylococcus aureus* is a Gram-positive pathogen responsible for the majority of skin and soft tissue infections (SSTIs). *S. aureus* colonizes the anterior nares of approximately 20-30% of the population and transiently colonizes the skin, thereby increasing the risk of developing SSTIs and more serious infections. Current laboratory models that mimic the skin surface environment are expensive, require substantial infrastructure, and limit the scope of bacterial physiology studies under human skin conditions. To overcome these limitations, we developed a cost-effective, open-source, chemically defined media recipe termed skin-like media (SLM) that incorporates key aspects of the human skin surface environment and supports growth of several Staphylococcal species. We utilized SLM to investigate the transcriptional response of methicillin-resistant *S. aureus* (MRSA) following growth in SLM compared to a commonly used laboratory media. Through RNA-seq analysis, we observed the upregulation of several virulence factors, including genes encoding functions involved in adhesion, proteolysis, and cytotoxicity. To further explore these findings, we conducted qRT-PCR experiments to determine the influence of media composition, pH, and temperature on the transcriptional response of key factors involved in adhesion and virulence. We also demonstrated that MRSA primed in SLM adhered better to human corneocytes and demonstrated adhesin-specific phenotypes that previously required genetic manipulation. These results support the potential utility of SLM as an *in vitro* model for assessing Staphylococcal physiology and metabolism on human skin.

**Importance:** *Staphylococcus aureus* is the major cause of skin diseases, and its increased prevalence in skin colonization and infections present a need to understand its physiology in this environment. The work presented here outlines *S. aureus* upregulation of colonization and virulence factors using a newly developed media that strives to replicate the human skin surface environment, and demonstrates roles for adhesins ClfA, SraP, and Fnbps in human corneocyte adherence.

## Introduction

The skin is the largest organ in the human body (1). This organ is highly stratified and interspersed with skin appendages including hair, sebaceous glands, eccrine glands, and apocrine glands (2). The stratum corneum is the outermost layer of the skin, and is comprised of 15 – 25 layers of nonviable, anucleate corneocytes that are assembled in a “brick and mortar” arrangement that prevents water loss and protects the body against environmental challenges and pathogens (2). Over the course of human evolution, encoded functions of the human skin have acquired several mutations that resulted in a unique skin surface environment encountered by commensals and pathogens alike (3). For example, the ubiquitously distributed eccrine glands produce sweat at up to 0.5 – 3.5 L/h combined and are a unique attribute of human skin physiology (3–5). Additionally, the human skin surface is more acidic (pH 4.1 – 5.8) than that of other mammals commonly used in skin infection models (6, 7).

*Staphylococcus aureus* is the most common pathogen isolated from skin and soft tissue infections (8). Treatment and prevention of *S. aureus* skin infections are further complicated by the prevalence and transmission of community-acquired, methicillin-resistant *S. aureus* (MRSA) strains (8–10). The CA-MRSA USA300 strain, in particular, has become epidemic and drives the increased prevalence of SSTIs in the last two decades (10). MRSA infection risk is correlated to nasal carriage, a reservoir from which MRSA can transiently colonize the skin (11).

Studies of Staphylococcal physiology on human skin in the early stages of colonization, and preceding infection, are restricted by cost and other limitations of currently used laboratory models (7, 12). To close this gap, we developed a skin-like media (SLM) that incorporates metabolites derived from eccrine sweat and stratum corneum turnover. We also incorporated the acidity, temperature, and buffering capacity of a healthy skin barrier into the media recipe and growth conditions. This media promoted growth of seven different Staphylococcal species isolated from both infectious and colonization contexts. To inquire how MRSA responds to these conditions, we analyzed transcriptional changes of the CA-MRSA USA300 strain LAC* following growth in SLM in comparison to a commonly used growth media, TSB. These transcriptional changes were compared to qRT-PCR data from previously published *ex vivo* human skin explant experiments as a comparator. Differentially regulated genes encoding key colonization and virulence factors were validated by qRT-PCR, and the influence of temperature and pH on the transcription of these loci was assessed. The utility of SLM was demonstrated with assessment of MRSA adhesion to human corneocytes, an assay that previously required regulatory manipulation of adhesins to assess their role in this key aspect of skin colonization. The results support that growth in SLM stimulates a dramatic transcriptional response in MRSA, including in the expression of key virulence factors, and can expand the applicability of host-microbe studies performed at the research bench.

## Results

### Considerations and development of the SLM

The skin-like media (SLM) formulation was informed by an extensive review of skin surface composition in the literature. Human sweat, produced by eccrine glands ubiquitously distributed on human skin, produce up to a combined 0.5 – 3.5 L of sweat every hour (Figure 1A) (4, 5). We incorporated the constituents of human sweat into the media, including Na^+^, K^+^, Cl^-^, lactate, urea, Mg^2+^, and Ca^2+^ (Figure 1B, Table S1). Additionally, we included amino acids derived from the natural moisturizing factor (NMF), a product of corneocyte turnover, in the form of water-soluble and DMSO-soluble 40X amino acid mixes (Figure 1B, Table S1) (13). This mixture included the skin-specific amino acid derivatives *trans-* urocanic acid and 2-pyrrolidone-5-carboxylic acid (14, 15). Essential vitamins, sodium phosphate, and glucose were also included to provide vital nutrients for Staphylococci (Figure 1B, Table S1). While glucose concentration on the skin surface is relatively unexplored, we added 1 mM glucose based on measurements of dermal tissue and eccrine sweat gland secretions (4, 16). The human-specific fatty acid, sapienic acid, was integrated to simulate fatty acid stress of the skin while maintaining the ability to collect absorbance-based culture density measurements (Figure 1, Table S1) (17, 18). The media was buffered to pH 4.8 using 100 mM MES buffer to replicate the skin’s pH-buffered nature (19). Lastly, we set the culturing temperature to 32 °C, mirroring the temperature of the anterior forearm (20). Component concentrations were determined using available quantitative data, and maintaining relative proportions when absolute quantifications were not available.

**Figure 1.**
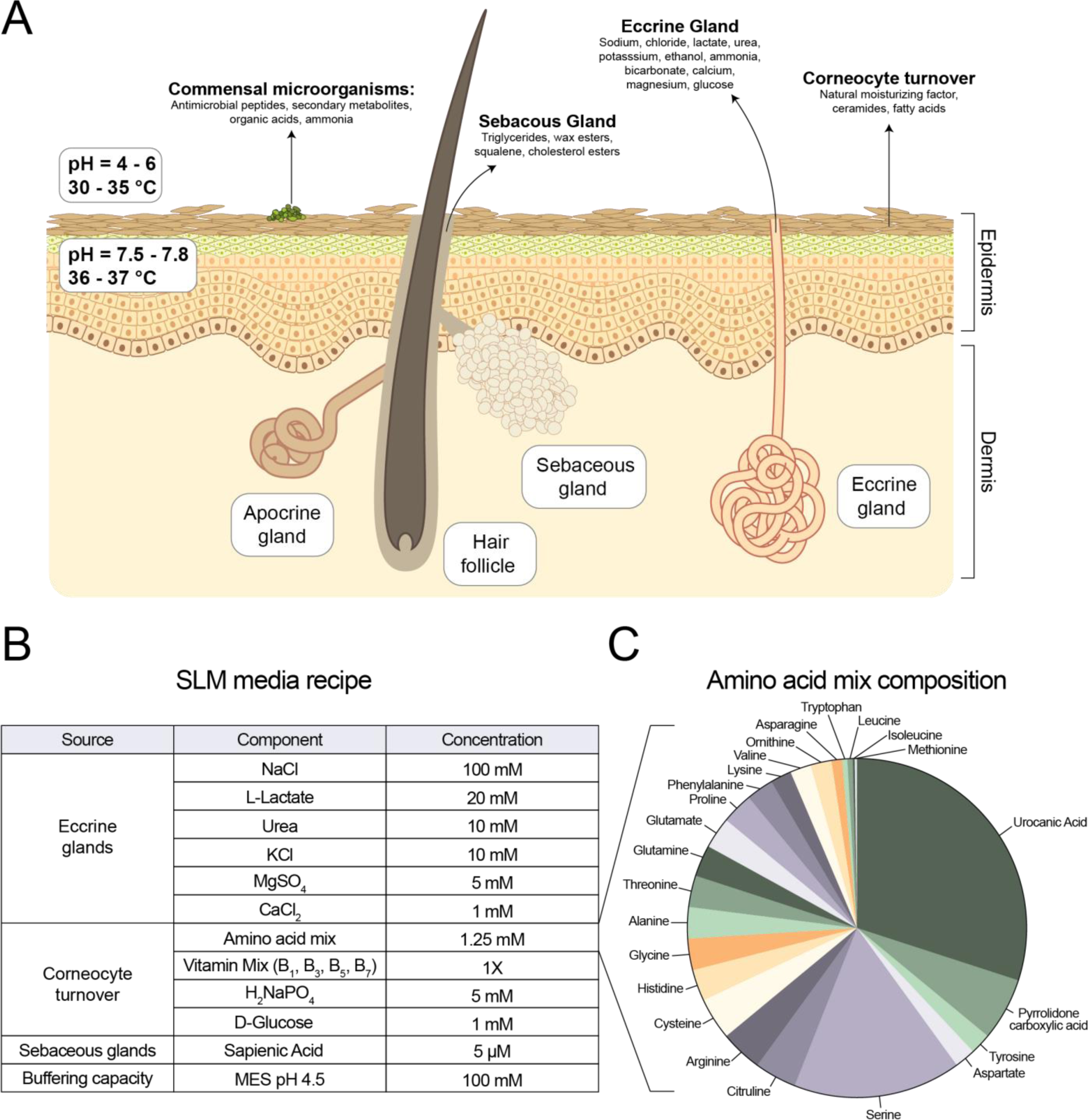
Development of the skin-like media (SLM) recipe. (A) Aspects of the human skin environment that were considered in the development of the media. (B) Components incorporated into the media recipe and the concentrations incorporated into the media. (C) A pie chart demonstrating the relative concentrations of amino acids by molarity incorporated into the SLM.

To gauge *S. aureus* growth response to the SLM composition, we tested different concentrations of lactate, urea, amino acid mix, glucose, sapienic acid, and buffer with *S. aureus* strain LAC*, a derivative of the epidemic MRSA strain USA300_LAC that has been cured of plasmid p03, hereafter referred to as MRSA (Figure S1) (21, 22). Lactate positively influenced growth within 10-30 mM, becoming growth inhibitory beyond 50 mM (Figure S1A). Similarly, urea’s growth-enhancing effect plateaued at 50 mM (Figure S1B). The amino acid mixture proved indispensable for growth, which was unsurprising since MRSA has documented amino acid auxotrophies (Figure S1C) (23). The amino acid mixture was inhibitory above 3.75 mM, likely due to the DMSO concentration required to keep amino acids such as *trans-*urocanic acid soluble (Figure S1C). Surprisingly, glucose did not improve growth beyond 1 mM (Figure S1D). Sapienic acid, which has documented inhibitory properties with Staphylococci, hindered growth of MRSA between 20-50 μM (Figure S1E). Finally, buffer concentration influenced growth rates, but not final yields (Figure S1F). Addition of trace metals did not affect growth and was not included in the final recipe (data not shown).

### Growth of diverse Staphylococci in SLM

We next tested growth of different Staphylococcal strains from infection and skin colonization contexts, including *S. aureus* and *S. epidermidis* isolates from healthy donors and atopic dermatitis patients (Table S2). All isolates demonstrated growth in SLM, with growth rates and yields varying across isolates within *S. aureus, S. epidermidis, S. hominis, S. capitis, S. lugdunensis, S. warneri,* and *S. haemolyticus* species (Figure 2). Variability appeared to be isolate-specific, and not applicable to an entire species.

**Figure 2.**
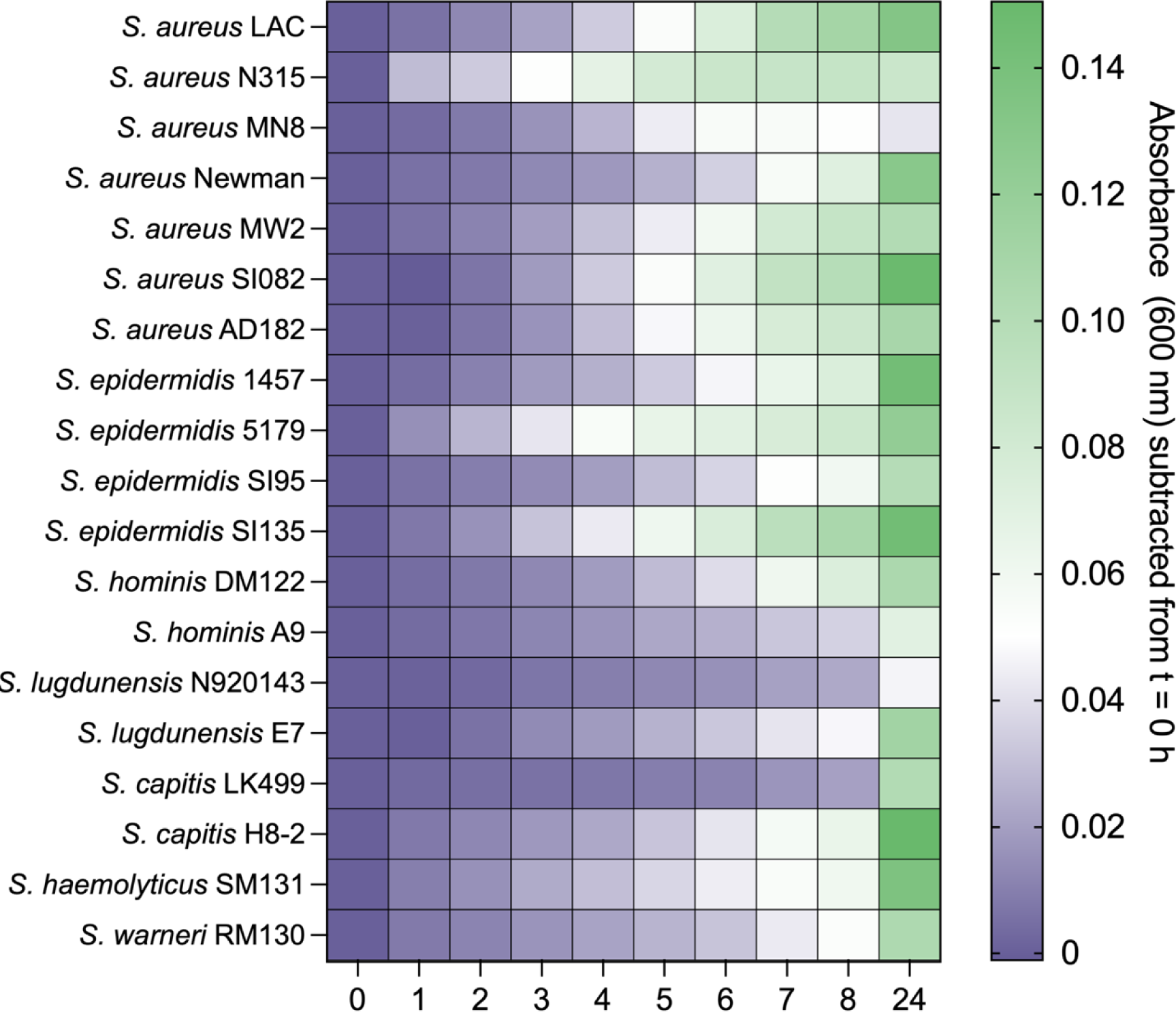
Growth of Staphylococcal isolates in SLM. Growth of isolates from varied infection/colonization contexts, strain backgrounds, and Staphylococcal species were tested in SLM. Heterogeneity in growth rates and growth yield after 24 hours did not appear to be species-dependent.

### Transcriptional response of MRSA in SLM

For insights into MRSA transcriptional response to SLM, we compared MRSA grown in SLM to cultures grown in TSB. Cultures were collected at the same culture density in mid to late exponential growth to normalize *agr*-dependent effects on transcription (Figure 3A) (24). A volcano plot generated from the RNA-seq differential expression analysis demonstrated that 1156 of the 2629 annotated gene loci were >2-fold differentially regulated and met the adjusted p-value cutoff (*p_adj_*<0.05, Figure 3B, Table S3). Analysis of the upregulated genes using ShinyGO pathway enrichment analysis (http://bioinformatics.sdstate.edu/go/) showed that top enriched pathways in upregulated genes included purine biosynthesis (*pur* genes), branched-chain amino acid biosynthesis (*ilv* and *leu* genes), nickel cation binding (*ure* operon), and extracellular proteases (*spl* and *ssp* genes) (Figure 3C). Top enriched pathways in downregulated genes included functions associated with transcription and translation machinery (polymerase, ribosomal, tRNA genes, rRNAs) and pyrimidine biosynthesis (*pyr* genes, Table S3, Figure 3D). The downregulated functions largely reflect the difference in growth rate between the two conditions.

**Figure 3.**
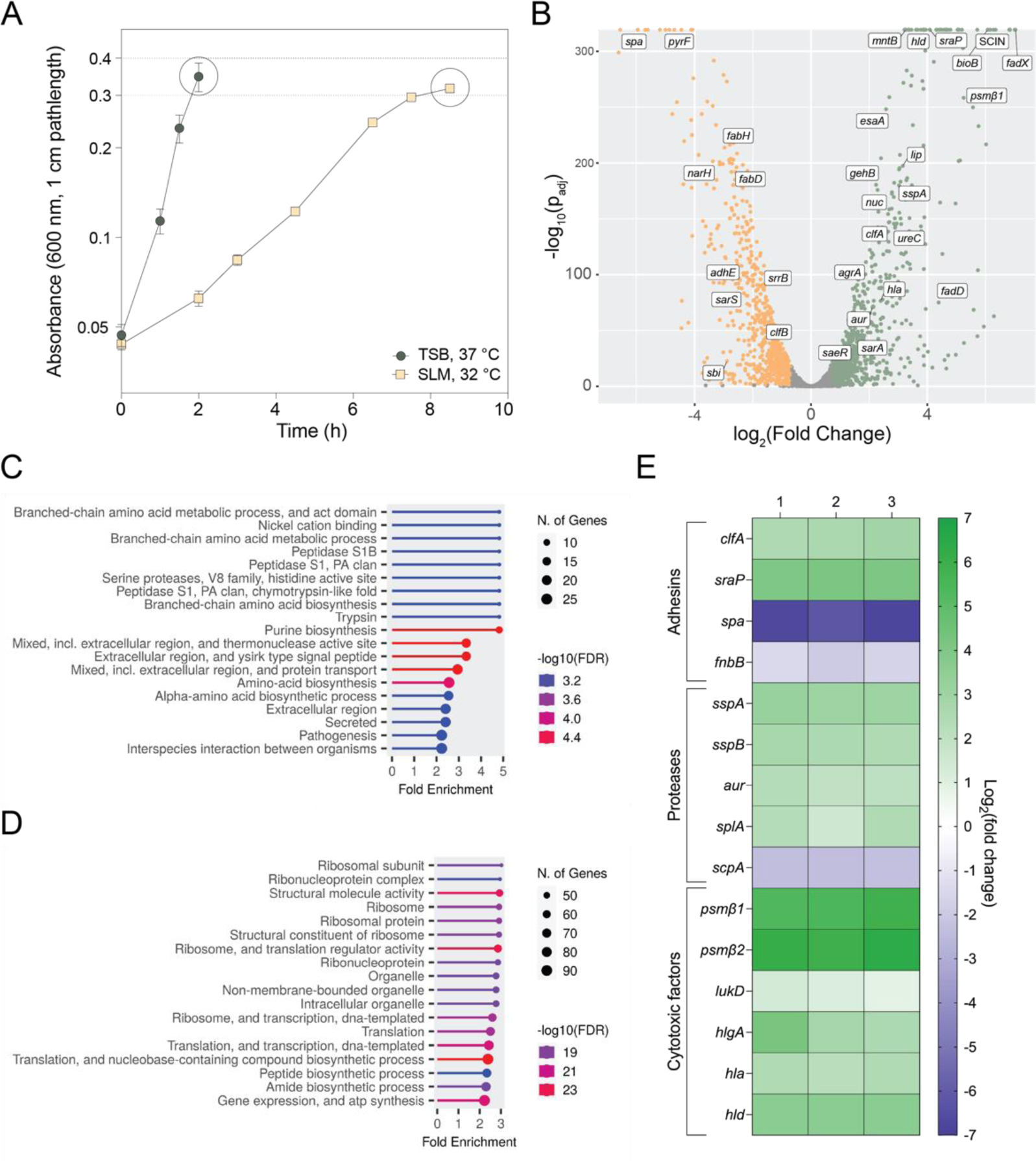
Transcriptional response of *S. aureus* to SLM. (A) Samples of *S. aureus* LAC* were collected at OD_600_ = 0.3 following growth in TSB at 37 °C or SLM at 32°C. (B) A volcano plot depicting the differential expression (x-axis) and adjusted p-value (y-axis) of each gene in the *S. aureus* LAC* genome. Upregulated genes are in green and downregulated genes are in orange. A subset of genes are labeled. (C) Enriched pathways in the upregulated genes. (D) Enriched pathways in the downregulated genes. (E) Log_2_(fold-change) of significantly differentially regulated adhesins, proteases, and cytolytic factors that play known roles in *S. aureus* colonization and virulence.

Of note, regulation of many colonization and virulence factors were differentially expressed in SLM compared to TSB (Figure 3B, 3E, Table S3). Upregulated adhesins included <u>cl</u>umping factor <u>A</u> (*clfA,* SAUSA300_0772) and <u>s</u>erine-rich <u>r</u>epeat glycoprotein <u>a</u>dhesin (*sraP* or *sasA*, SAUSA300_2589). A few adhesins were downregulated, including surface protein A (*spa*, SAUSA300_0113) and fibronectin-binding protein B (*fnbB,* SAUSA300_2440). All extracellular proteases (*splA-F,* SAUSA300_1758-1753; *aur,* SAUSA300_2572; *sspA-B,* SAUSA300_0950-1) with exception of staphopain A (*scpA*, SAUSA300_1890), were upregulated (Figure 3E, Table S3). Several cytolytic factors and toxins were also upregulated. Most notable were the upregulation of phenol soluble modulins (PSM) β1 and β2 (*psmβ1-2,* SAUSA300_1067-1068), which were among the most highly upregulated genes in the dataset. Other cytolysins such as leukotoxins (*lukD,* SAUSA300_1768), and hemolysins (*hla,* SAUSA300_1058; *hlgABC,* SAUSA300_2365-2367; *hld,* SAUSA300_1988) were significantly upregulated in the RNA-seq dataset (Figure 3E, Table S3). Collectively, these changes suggest that MRSA adapts to a skin-like environment by increasing expression of virulence factors and metabolic pathways like the purine and amino acid biosynthesic pathways.

### RNA-seq comparison to MRSA transcriptional response to an *ex vivo* model

Although comparison of the SLM RNA-seq to an *in vivo* RNA-seq dataset was not possible, we found transcriptional changes in colonization and virulence factors following growth in SLM aligned with published qRT-PCR data following MRSA application on human skin explants for 24 and 72 hours (Figure S2, Table S4). Remarkably, none of the loci tested in the explant study conflicted with transcriptional responses in SLM and human skin explants (Figure S2) (25). Shared upregulated genes across all comparisons included *asp23, atl, aur, clfA, ebpS, esaA, essB, esxA-C, mntA,* SAUSA300_0883, *sraP, sspA,* and *sspB* (Table S4). Shared downregulated genes included *clfB, sarS,* and *spa* (Table S4). These results suggest that MRSA grown in SLM shares some common regulatory responses with the human skin surface.

### Influence of pH and temperature on MRSA virulence factor expression

To validate reproducibility of the transcriptional changes in the RNAseq and examine the impact of temperature and pH on regulation, cultures were grown in TSB and SLM at both 32 and 37 °C, and at pH 4.8 and pH 7.2. Purified RNA was converted to cDNA, and transcription of adhesins (*sasG, clfA, sraP, spa, fnbA, fnbB*), proteases (*sspA, aur, splA, scpA*), and cytolytic factors (*psmα, psmβ, hlgA, lukS, hla*) were quantified by qRT-PCR using the primers listed in Table S5, and the C_t_ values were compared to TSB at 37 °C and neutral pH (Figure 4). MRSA grown in SLM and TSB at both 32 and 37 °C showed some differences in transcription due to temperature (Figure 4A). Principal component analyses of these data showed separation of TSB and SLM samples along PC1, suggesting that media composition and/or pH exerted a stronger influence than temperature on the regulation of these factors (Figure 4B). Temperature also appeared to exert a stronger influence on regulation of these factors in SLM than in TSB, as noted by the separation of the SLM samples but not the TSB samples at the two temperatures (Figure 4B). The same approach was repeated at both neutral (pH = 7.2) and acidic (pH = 4.8) conditions (Figure 4C-D). These data showed more dramatic pH-dependent changes in transcription (Figure 4C). PCA of these data showed the confidence intervals of SLM and TSB samples at pH 7.2 overlapped (Figure 4D). When SLM and TSB media were at pH 4.8, they separated from the neutral media conditions along the first and second components, respectively (Figure 4D). These data suggest that pH and temperature have important roles in transcription of MRSA virulence factors, and media composition influences the regulatory outcome of these signals.

**Figure 4.**
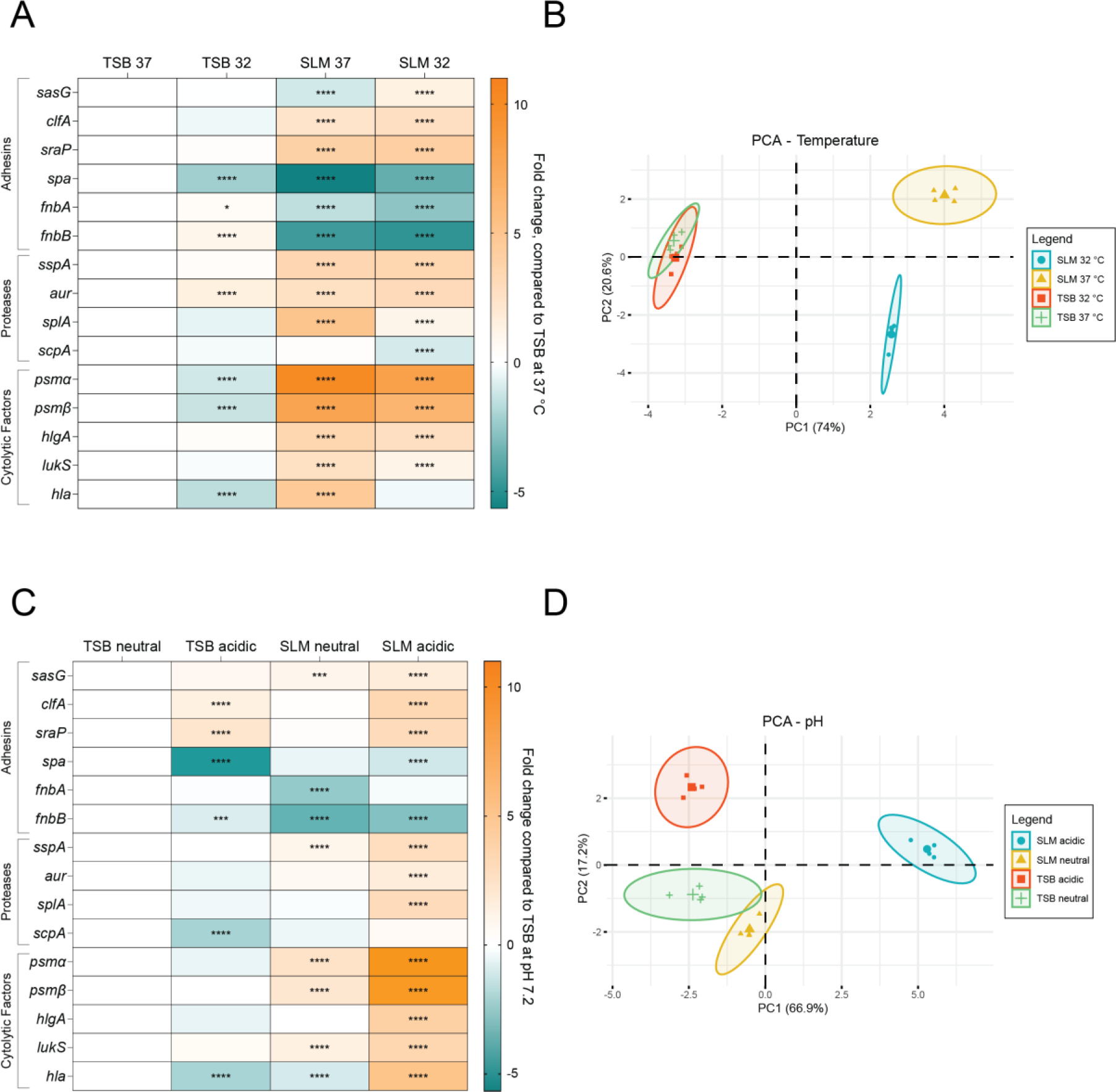
Effect of temperature and pH on virulence factor expression in TSB and SLM. A) qRT-PCR in TSB and SLM at 32 °C and 37 °C of transcripts of genes encoding key virulence factors. B) PCA plot of the fold-changes shown in A, color-coded by condition. C) qRT-PCR in TSB and SLM at pH 7.2 (neutral) and pH 4.8 (acidic) of transcripts of genes encoding key virulence factors. D) PCA plot of the fold-changes shown in Figure 6, color-coded by condition. Significant fold-changes were determined using a two-way ANOVA and are denoted with asterisks; * p<0.05, ** p<0.01, *** p<0.001, **** p<0.0001

### Enhanced Adherence of MRSA to Human Corneocytes following growth in SLM

Previously, studies of MRSA adherence to human corneoocytes required deletion of the regulator MgrA (Δ*mgrA*) or overexpression constructs to assess an adhesin’s role in corneocyte adherence (26, 27). Since genes encoding several adhesins including CflA and SraP were significantly upregulated in SLM in the qRT-PCR analysis, we tested whether MRSA grown in SLM adhered better to human corneocytes from the stratum corneum than when grown in TSB. With exception of the comparison in Figure 5A, MRSA adherence to corneocytes significantly increased following growth in SLM compared to TSB (Figure 5). The improved adherence in SLM was abrogated by mutation of *clfA, fnbAB*, or *sraP,* suggesting that these adhesins all contribute to MRSA adherence to human corneocytes (Figure 5A-C). Deletion of *sasG* did not abrogate adherence (Figure 5D). Although baseline adherence varied from donor to donor, the trends across strains were consistent (Figure S3). Each input was dilution plated and enumerated for CFU/mL to ensure that inoculums were consistent across strains and culture conditions (Figure S4).

**Figure 5.**
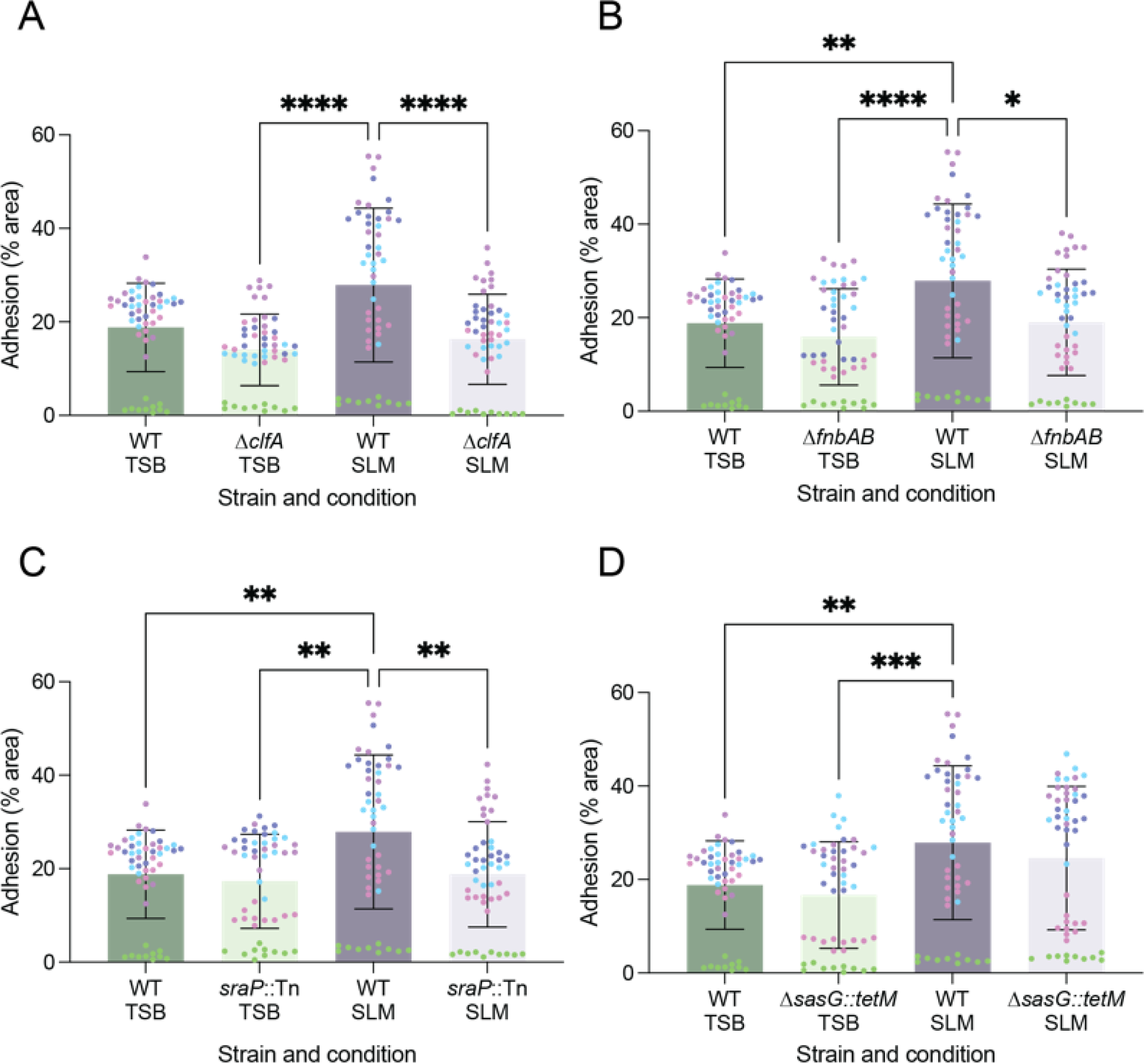
MRSA corneocyte adhesion increases following growth in SLM, and is dependent on ClfA, SraP, and Fnbs. Corneocyte adhesion assays were performed with cultures grown in TSB or SLM to OD_600_ ∼0.2. Cultures were assessed for adherence to corneocytes from five human donors (color-coded). The contributions of adhesins to corneocyte adhesion was assessed for (A) ClfA, (B) FnbA/B, (C) SraP, and (D) SasG. Cultures grown in TSB had no significant difference in adhesion. Adhesion of the WT was significantly higher in SLM compared to TSB in three of the comparisons (B-D), and ClfA, FnbA/B, and SraP contributed to adhesion in the SLM-primed cells (A-C). SasG did not contribute to adhesion, likely due to the point mutation in *S. aureus* LAC* that prevents anchoring of SasG to the cell wall. Statistics: Kruskal-Wallis test, p<0.05.

## Discussion

Research on bacterial behavior at the human skin barrier is hindered by the limitations of available models and the institutional infrastructure required for their utilization. In this study, we generated a skin-like media (SLM) recipe which replicates the acidity, ionic strength, buffering capacity, and known metabolic composition of an intact human skin barrier. This formulation was intentionally designed for easy assembly, requiring approximately 30 minutes once the necessary stocks are prepared. Moreover, the composition and pH can be adjusted for the investigation of skin niches, skin disorders, and particular metabolites of interest. Furthermore, the recipe is openly accessible for improvement as more insights into the skin’s surface environment become available. We note that another skin-like media recipe has been recently published (28). In the recipe reported herein, we have improved the incorporation of unique, skin-specific metabolites by recapitulating the measured amino acid composition of the natural moisturizing factor, which has not been included in the prior recipe.

We evaluated the growth of several Staphylococcal isolates from both infection and skin colonization contexts using this media. Overall, growth yields in SLM were lower compared to nutrient-rich TSB, which was expected given the relatively nutrient-scarce nature of the skin environment (29). All the Staphylococcal isolates tested grew in SLM, however, the growth rates and yields varied considerably across isolates. The observed variability could point to differences in the metabolic networks in these isolates, including isolate-specific auxotrophies that have not yet been identified. Exploring these metabolic differences among isolates and species in the future will contribute to a deeper understanding of the metabolic requirements of these microorganisms in their natural habitat.

The MRSA strain used in these studies was isolated from a skin abscess during an outbreak investigation and is utilized as a model strain to understand MRSA pathogenesis on the skin (30). As colonization often precedes infection, we aimed to investigate MRSA responses when encountering a skin-like environment (31, 32). Analysis of the transcriptional response of MRSA following growth in SLM, compared to TSB, revealed significant changes affecting a substantial portion of the genome. There were shifts in several metabolic pathways, including the upregulation of purine and branched-chain amino acid (BCAA) biosynthesis. Upregulation of purine biosynthesis was previously shown to be important for persistent infections (33), while upregulation of BCAA biosynthesis was accompanied by downregulation of the negative regulator, CodY, in SLM (34). The BCAAs are limited in abundance relative to other amino acids in the natural moisturizing factor, so it’s possible that MRSA initially relies on BCAA biosynthesis rather than scavenging (13). Most downregulated pathways identified in the pathway analysis involved transcription and translation machinery, however pyrimidine biosynthesis was identified as the most downregulated metabolic pathway in the RNA-seq dataset. It’s possible this transcriptional response prevents diversion of the intermediate phosphoribosyl pyrophosphate (PRPP) from purine biosynthesis.

Many upregulated genes encoded for secreted and cell wall-associated factors, including adhesins, extracellular proteases, and toxins. Genes encoding adhesins ClfA, SraP, and Ebh were upregulated in SLM. In contrast, the genes encoding IgG-binding adhesins Spa and Sbi were downregulated in SLM, as well as those encoding for adhesins ClfB and FnbB. The downregulation of FnbB was surprising, given its previously demonstrated role in corneocyte adhesion (27). In this work, we confirmed the role of the fibronectin-binding proteins in stratum corneum adherence and demonstrated roles for ClfA and SraP. The genes encoding extracellular proteases, with the exception of ScpA, were also upregulated. The downregulation of *scpA* in SLM may correspond to a healthy skin barrier, as qRT-PCR of mRNA collected from donors’ skin showed low expression of *scpA* in healthy controls compared to atopic dermatitis lesions, and compared to expression of *sspB* in both conditions (35). Significant progress has been made in characterizing the biochemistry and substrate affinity of the Spl proteases A-F (36–38). However, with exception of SplA, the roles of the Spl proteases in colonization or infection have not been established (39). Aureolysin initiates the proteolytic cascade that activates V8, which is involved in skin pruritis and in turn activates Staphopain B (40–42). Additionally, several cytolytic factors and toxins were upregulated. Most notable was the upregulation of PSMs (*psmα, psmβ1-2*) and other cytolysins such as leukotoxins (*lukD,* SAUSA300_1768) and hemolysins (*hla, hlgABC, hld*). PSMα and alpha toxin (Hla) both have roles in virulence and keratinocyte damage (43). Both lukS-PV and lukF-PV, which play critical roles in skin infection, had higher levels of expression in SLM compared to TSB (44). Athough the increase did not meet our 2-fold cutoff in the RNA-seq, we observed significant upregulation of *lukS* in the qRT-PCR experiments.

The observed upregulation of the PSMs, toxins, and extracellular proteases could be explained by the upregulation of Agr quorum sensing system, encoded by *agrBDCA* (the *agr* operon). The activation of Agr has previously been shown to be critical for MRSA-induced skin barrier damage and skin infections, and this damage can be mitigated by Agr inhibitors (45, 46). Although cultures were taken at the same density, the *agr* operon was upregulated ∼2-3-fold. Additionally, upregulation of delta toxin, encoded by *hld,* and downregulation of *spa* and its transcriptional activator, *sarS*, are consistent with activation of Agr (47, 48). Induction of Agr could possibly be explained by de-repression from σ^B^, the transcription of which was downregulated in SLM (49). These data support the critical role of the Agr regulon in colonization and virulence on the skin (50).

In the absence of an appropriate *in vivo* comparator for SLM, we compared our RNA-sequencing data with available qRT-PCR data obtained of the same MRSA strain grown on human skin explants for 2-3 days. Strikingly, the RNA-seq data of the tested loci aligned with the transcriptional responses observed in the qRT-PCR dataset, suggesting that for a subset of loci, *S. aureus* responds to the media environment similarly to how it would on the human skin surface. Most notable in this comparison was the upregulation of *clfA*. ClfA, an adhesin unique to the *Staphylococcus aureus* species, was also upregulated in another human skin explant dataset (25, 51).

Both the pH and temperature of the skin environment differ from the conditions typically used for Staphylococcal research *in vitro*. To understand how these factors influence regulation of MRSA colonization and virulence factors, we investigated the impact of temperature and pH on the transcription of 15 factors in both TSB and SLM. Although both parameters influenced the transcriptional response of the loci, the direction of the response was not always consistent. For instance, the expression of PSMs (*psmα, psmβ*) and alpha toxin (*hla*) displayed different responses to a change in pH in SLM and TSB. This discordance in response suggests that studying the effect of the skin-like environment on *S. aureus* regulation of these factors could offer insights into their roles in *S. aureus* skin colonization and virulence.

The significant shifts in regulation of the adhesins prompted investigation into *S. aureus* adherence to human corneocytes. While previous studies required genetic modifications to study *S. aureus* adherence to corneocytes (26, 27), we found that MRSA primed in SLM exhibited enhanced adherence to human corneocytes compared to priming in TSB. Additionally, adhesin phenotypes emerged in SLM-primed cultures that weren’t previously observed in TSB-primed cultures. In SLM-primed cultures, we identified the involvement of ClfA, the fibronectin-binding proteins (Fbps), and SraP in corneocyte adherence. While the roles of Fbps aligned with earlier research, the roles of ClfA and SraP in corneocyte adherence were novel findings (27). ClfA is unique to *Staphylococcus aureus,* and the consistent upregulation of ClfA in two *ex vivo* skin explant experiments and SLM, as well as its contribution to corneocyte adherence, point to a critical role for ClfA in initial skin colonization (25, 51). Of the adhesins tested, SasG was the only one that did not contribute to corneocyte adherence. This is likely because SasG is truncated in USA300 MRSA strains such as *S. aureus* LAC* (52). Baseline corneocyte adherence was much higher in these experiments than previously reported (26). To accommodate the growth yields in SLM, cultures were collected in TSB and SLM at a lower culture density than previously used. Additionally, significant differences in adherence were observed across donors used in this study, which may point to host factors also playing a role in bacterial adherence.

It’s important to note that this study has limitations, such as the limited availability of comprehensive, quantitative metabolite analyses of the skin surface to inform the recipe design, and the absence of publicly available RNA-sequencing analyses of *S. aureus* on an *ex vivo* human skin model. As more information becomes available, the media recipe’s composition can be further refined, as has been observed in the iterative development of synthetic sputum media (53). Additionally, only the growth of Staphylococci was tested in this media. The study of organisms such as *Corynebacterium*, which rely on lipids as a key carbon source, would likely require addition of a synthetic sebum as described previously (28).

The aim of this work was to develop a skin-like media recipe that facilitates cost-effective and high-throughput study of Staphylococcal metabolism and physiology on the skin surface. The transcriptional response of MRSA in SLM aligns with publicly available qRT-PCR analyses of the same MRSA strain on human skin explants. Furthermore, priming MRSA strains in SLM enabled the examination of adhesin contributions to corneocyte adherence without regulatory manipulation. Collectively, this work introduces another avenue for studying Staphylococcal physiology within the context of the skin environment.

## Methods

Strain construction. The Δ*sasG::tetM* deletion construct was inserted into pJB38 using EcoRI and SalI restriction cloning sites, comprising of an fragment upstream of *sasG* amplified with primers HC246 (ATGGAATTCAATGATTTGAAAAGCAAGAGCAATA) and HC247 (CTCGAGGGTACCGCTAGCATCTCTCATTTGCATACTCCTTTTTCC), and a fragment downstream of *sasG* amplified with primers HC248 (GCTAGCGGTACCCTCGAGGCTGGATTAATGTTATTGGCACGT) and HC249 (GATGTCGACTTGATGTTATTGCAAGTAAAGGAAT). The tetracycline resistance marker *tetM* was amplified using primers HC3 (GTTAGCTAGCCCTAGGCAAATATGCTCTTACGTGC) and HC4 (GGCATGCTAGCGCACTAAGTTATTTTATTGAACATATATCTTAC) and inserted into the NheI site included between the upstream and downstream fragments by the above primers. The resulting vector, pHC128, was used to generate the *sasG::tetM* deletion in LAC as described previously (54).

### Growth media

Trypto soy broth (TSB, RPI Cat. No. T48500) at 30 g/L was used for overnight cultures and as the reference culture conditions for transcriptional analyses and phenotypic assays. For pH studies, TSB was acidified to pH 4.8 using HCl. Skin-like media (SLM) was assembled fresh using the prepared stocks and recipe outlined in Table S1A.

### Preparation of SLM

Stocks of NaCl, L-lactate, urea, KCl, H_2_NaPO_4_, MgSO_4_, CaCl_2_, and D-glucose were prepared in water and filter sterilized with a 0.22 µm MCE filter (Millipore Sigma Cat. No. GSWP04700, Table S1A). The MES stock was filter sterilized using a 0.22 µm PES filter (VWR, Cat. No. 73520-982). The 1000X vitamin mix stock was prepared in water using the recipe outlined in Table S1B and filter sterilized with a 0.22 µm MCE filter. The 40X amino acid mix was separated into DMSO-soluble (Table S1C) and water-soluble (Table S1D) mixtures. These stocks were generated as dry mixes by weighing out the grams indicated in the “g/L” column and blending into a fine powder using a mortar and pestle. The dry mixes were stored in falcon tubes in a desiccating chamber at 4 °C. The 40X DMSO-soluble amino acid mix was weighed out fresh for each experiment, solubilized in DMSO at 2.81 mg/mL, and filter-sterilized with a 0.22 µm nylon filter (VWR, Cat. No. 7649-030). The 40X water-soluble amino acid mix was also weighed out fresh for each experiment, solubilized in water at 4.10 mg/mL, and filter-sterilized with a 0.22 µm MCE filter. For the qRT-PCR pH studies, 1M MOPS pH 7.4 stock was prepared and replaced the 1 M MES pH 4.5 stock in the media recipe. <u>Growth conditions</u>. Strains were grown in biological triplicate or quadruplicate in 5 mL of TSB shaking at 250 RPM in 18 mm x 150 mm glass culture tubes at 37 °C for 16 hours. For RNA-sequencing and qRT-PCR, overnight cultures were subcultured at 1:100 (v/v) dilution into flasks containing TSB or SLM (20% media volume:flask volume). Cultures were shaken at 250 RPM at 32 °C for SLM and 37 °C in TSB to an OD of 0.2-0.3, as measured at 600 nm with a 1 cm pathlength cuvette. For growth measurements of *S. aureus* LAC* and other Staphylococci as shown in Figure 2 and Figure S1, cultures were subcultured at 1:100 (v/v) dilution in a total volume of 150 µL SLM in 96-well plates, and each biological replicate was plated in technical triplicate (Corning 3370, Fisher Scientific Cat. No. 07-200-656). Cultures were incubated at 32 °C at 1000 RPM (Stuart SI600 Incubator, Cole-Palmer) and at each desired timepoint, culture density was measured by absorbance at 600 nm (Tecan Infinite200-PRO, Tecan). Technical replicates were averaged together at each timepoint for data analysis.

### RNA purification

Cultures were collected and centrifuged in falcon tubes at 3,900 *xg* for 10 min. Supernatant was discarded, and the cell pellets were quickly resuspended in 1 mL ice-cold 1X PBS and moved to a 1.7 mL microcentrifuge tube. Tubes were centrifuged at 16,000 *xg* for 30 seconds. Supernatant was aspirated, and cell pellets were flash-frozen in dry ice before storage at -80 °C. RNA purification was performed within 2 weeks of freezing cell pellets as described previously (55).

### RNA-sequencing and data analysis

Purified RNA samples were submitted for rRNA depletion and mRNA sequencing (SeqCenter, Pittsburgh, PA). Samples were DNAse treated with Invitrogen DNAse (RNAse free). Library was prepared using Illumina’s Stranded Total RNA Prep Ligation with Ribo-Zero Plus kit and 10bp IDT for Illumina indices. Sequencing was done on a NextSeq2000 giving 2×51bp reads, 12M paired-end reads. Demultiplexing, quality control, and adapter trimming was performed with bcl-convert (v3.9.3) Quality of resulting reads were checked using FastQC (56). Adapter removal and read trimming was performed using Trimmomatic, using the paired-end option with a minimum read length of 36 nucleotides and trimming quality of 3 (57). After trimming, read quality was again assessed with FastQC. Trimmed reads were aligned to the *S. aureus* subsp. aureus USA300_FPR3757 reference genome (taxonomy ID: 451515) and read counts quantified using EDGE-pro (58). Un-normalized read counts were imported into R-Studio, and differential expression between SLM and TSB samples was quantified using DESeq2 with normalization (59). The volcano plot shown in Figure 3B was also generated in R-Studio using the ggplot2 package.

### qRT-PCR and data analysis

Purified RNA was treated with DNase per manufacturer’s instructions (Turbo DNA-*free* Kit, ThermoFisher Scientific, Cat. No. AM1907). DNase-treated RNA was quantified with a Qubit 4 Fluorometer and the Qubit RNA Broad Range Assay Kit per manufacturer’s instructions (ThermoFisher Scientific, Cat. No. Q10211). 1 µg of RNA was then converted to cDNA per manufacturer’s instructions and diluted to a working stock of 5 ng/µL (SuperScript IV VILO Master Mix, ThermoFisher Scientific, Cat. No. 11756050). qRT-PCR primers were designed using Primer 3 software (https://primer3.ut.ee/) and validated using a primer efficiency cutoff of >95%. qRT-PCR targets and associated qRT-PCR primers are listed in Table S5. Quantitative PCR was performed using PowerTrack SYBR Green Master Mix (ThermoFisher Scientific, Cat. No. A46109) and fluorescent intensity was monitored after each cycle (Bio-Rad CFX96, Bio-Rad). Reverse transcriptase-negative cDNA samples were included as a DNA contamination control. C_t_ values were obtained using the CFX Maestro Software (Bio-Rad) and fold-changes were calculated for each locus using the 2^-ΔΔCt^ method and *gyrB* as the reference gene (60, 61).

### Corneocyte adherence assay

Corneocytes were collected as described previously from five healthy donors under IRB protocol 19-2218 with their informed consent (26). Bacterial strains containing the GFP-expressing vector pCM29 were used and are listed in Table S2. Strains were grown in 5 mL TSB supplemented with 10 µg/mL chloramphenicol for 16 h at 37 °C and 250 RPM and subcultured at a 1:100 (v/v) dilution into flasks containing TSB or SLM (20% media volume: flask volume) supplemented with 2 µg/mL chloramphenicol. Cultures were grown at 37 °C for TSB and 32 °C for SLM to an OD_600_ 0.2-0.25 as measured by cuvette (1 cm pathlength). Cultures were centrifuged at 3,900 *xg* for 10 minutes at ambient temperature, and supernatant was aspirated. Cells were washed once in 1X PBS and normalized to ∼5 x 10^7^ CFU/mL in 1X PBS. Corneocyte adherence assay and data analysis were completed as previously described (26).

## Data availability statement

RNA-sequencing data were deposited in the NIH Sequence Read Archive (SRA) under BioProject PRJNA1029187.

## Acknowledgements

This work was funded by the NIH/NIAID grant(s) AI153185 and AI083211 to A.R.H., VA Merit Award BX002711 to A.R.H, and NIH/NIAID F32 fellowship AI172080 to F.G.C.

